# Meaning-making behavior in a small-brained hominin, *Homo naledi*, from the late Pleistocene: contexts and evolutionary implications

**DOI:** 10.1101/2023.06.01.543135

**Authors:** Agustin Fuentes, Marc Kissel, Penny Spikins, Keneiloe Molopyane, John Hawks, Lee R. Berger

## Abstract

Explorations in the Dinaledi Subsystem of the Rising Star cave system have yielded some of the earliest evidence of a mortuary practice in hominins. Because the evidence is attributable to the small-brained *Homo naledi*, these analyses call into question several assumptions about behavioral and cognitive evolution in Pleistocene hominins. The evidence from the Dinaledi Subsystem, and at other locations across the Rising Star cave system may widen the phylogenetic breadth of mortuary, and possibly funerary, behaviors. These discoveries may also associate the creation of meaning making and increased behavioral complexity with a small-brained hominin species, challenging certain assertions about the role of encephalization and cognition in hominin and human evolution. We suggest that the hominin socio-cognitive niche is more diverse than previously thought. If true, technological, meaning-making activities, and cognitive advances in human evolution are not associated solely with the evolution of larger brained members of the genus *Homo*.

**One-Sentence Summary:** Evidence for complex behaviors associated with a small-brained hominin suggest that large brains are not solely responsible for the manifestation of human-like behavioral complexity.

---

Contemporary humans engage in shared meaning via vocal, visual, tactile, and scent communication, often involving patterned use of bodies, objects, and materials. Of the very broad range of behaviors that create embedded shared meaning, the only ones that persist in archaeological contexts are those that leave material traces. Examples include aspects of mortuary behavior, engravings or paintings, personal ornaments, and use of ochre or other pigments. Prior to this century, most researchers readily accepted that such material traces could be interpreted as products of meaning-making behaviors when associated with *Homo sapiens*. But when such traces were associated with Neanderthals or other members of the genus *Homo* it was controversial. Previously, some archaeologists argued that the spectrum of behaviors in contemporary humans that involve shared meaning emerged as an integrated package in Late Pleistocene Africa, likely related to the dispersal of “modern” *Homo sapiens* throughout the world (e.g., Klein 1995). This view gave way to a recognition that meaning-laden material was used across a wider span of time, at least across part of the Middle Pleistocene (McBrearty and Brooks 2000), and not only by recent humans. In the last two decades, substantial evidence emerged of the extent of material evidence of meaning-laden behavior attributed to Neanderthals and other members of the genus *Homo* (Kissel and Fuentes 2018, 2021; Meneganzin and Killin 2024).

Currently a broad set of data demonstrates that some of these complex behaviors that involve shared meaning were manifested by multiple species and populations of the genus *Homo* including *Homo heidelbergensis, Homo erectus,* and possibly others (Galway-Witham, Cole, and Stringer 2019, Kissel and Fuentes 2018, 2021; Scerri and Will 2023) (**Figure 1).** Engravings of shell, bone, or rock surfaces have been identified in Middle Pleistocene or earlier contexts far from a range of African *Homo* populations. Some are also likely associated with *Homo erectus* (Joordens et al. 2014) as well as hominin populations that may have predated early Neanderthals in what is now Europe (Mania and Mania 1988; Sirakov et al. 2010). Evidence of ochre use occurs in archeological contexts across Africa and the Levant prior to 350,000 years ago, a time when *H. sapiens* has not yet been identified there (Ronen et al. 1998; Watts, Chazan, and Wilkins 2016; Dapschauskas et al. 2022). The control of fire by hominins is demonstrated in Early and Middle Pleistocene contexts where most researchers accept that *H. erectus* was present (Brain and Sillen 1988; Alperson-Afil 2008; Goren-Inbar et al. 2004; Hlubik et al. 2019; MacDonald et al. 2021). Mortuary evidences are claimed in association with hominins that predate or are not *H. sapiens* (Carbonell and Mosquera 2006). These geographically and temporally varied instances could be the result of taphonomic or dating issues (Püschel et al. 2021), but it is likely, given the increasing diverse temporal and geographic discovery of these behaviors and material that such complex behaviors associated with shared meaning were manifested by multiple populations/species of the genus *Homo* in addition to *Homo sapiens*. But how important was brain size to the evolution of these behaviors?

**Fig. 1.**
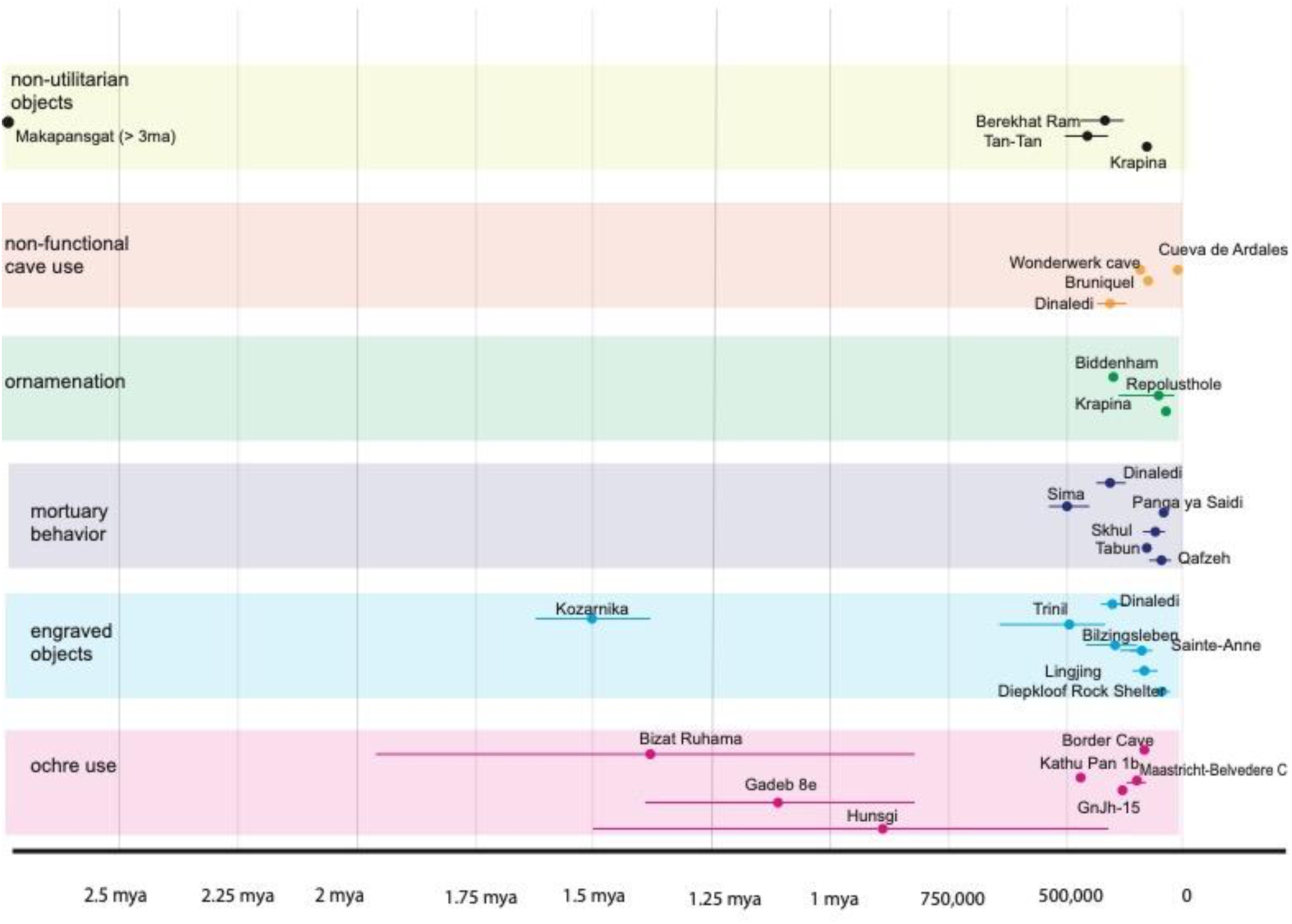
Archaeological evidence of culturally-mediated, meaning-making, behaviors. Dots represent different sites and the error bars are the maximum and minimum dates when available. See Table for details and See supplemental material for references.

There is substantive evidence that approximately 250-350,000 years ago *Homo naledi*, a small-brained hominin, transported deceased conspecifics into difficult to access locations in the Rising Star Cave system in what would, in humans, be described as a mortuary behavior (Berger et al. 2024). The use of deep areas of the Rising Star cave system for these behaviors implies considerable social collaboration, coordination, and planning. In the context of the subterranean Dinaledi Subsystem, these activities likely also required a light source, again implicating a depth of planning and coordination. What stands out as a possible contradiction is that *Homo naledi* fossil crania are small. With endocranial volumes ranging between 450 ml and 610 ml, this species overlaps in brain size with australopithecines, having smaller brains on average than *Homo erectus* and much smaller than modern humans or Neanderthals (See Figure 2 and section Reconsidering Brain Size).

**Fig. 2.**
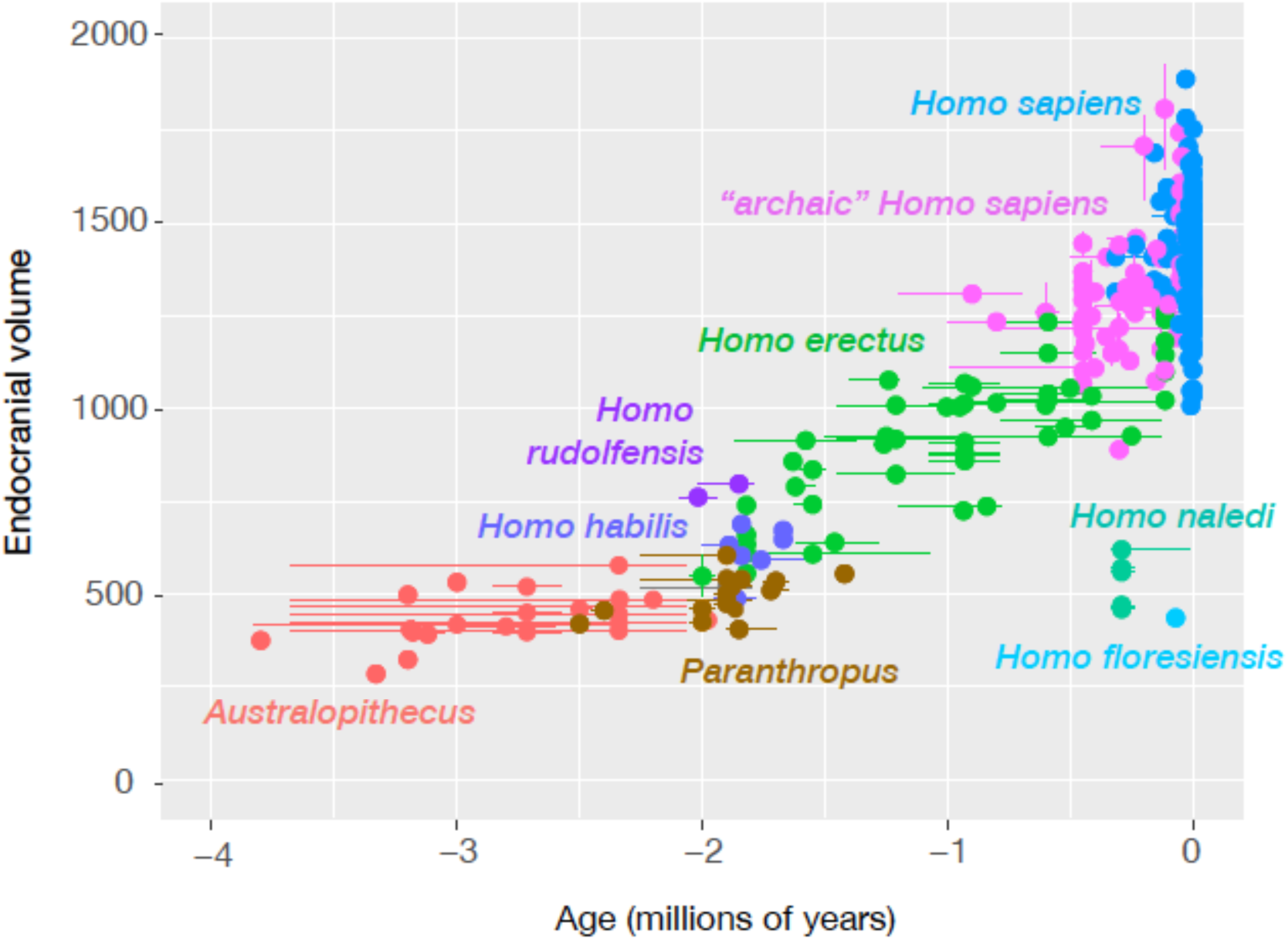
Endocranial volume estimates for hominin cranium. Error bars represent the maximum and minimum ages for specimens when available. See supplemental material for references. Hawks, John. 2023. Endocranial volumes for fossil hominins (dataset). Figshare https://doi.org/10.6084/m9.figshare.22743980

The information from *H. naledi* cannot be considered in isolation; it joins the broad array of data for meaning-making in Pleistocene hominins (Kissel and Fuentes 2018, 2021; Malfouris 2013). These behaviors reflect social groups that maintained solidarity, social coordination and cooperation in a mode not evident in living great apes but characteristic of contemporary humans. Here we offer an analysis of the reported complex behavior in the small-brained *Homo naledi* and suggest a suite of implications this has for our understanding of the relationships between brain size, cognition, complex behavior and the evolution of the genus *Homo* across the Pleistocene. These implications also query the driving forces behind encephalization and its relationship to the emergence of complex behaviors in hominins and other animals (see Tatersall 2023).

## Mortuary behavior as meaning making

*Mortuary behavior* has been defined as actions by individuals relating to the death of other individuals. Many kinds of non-human animals have been observed to engage in mortuary behavior upon the death of another individual in the same social group (Piel and Stewart 2015; Gonçalves and Carvalho 2019; Pettitt and Anderson 2020). Mortuary behavior may include the manipulation, inspection, or movement of dead bodies or body parts, or modifying behavior in proximity to dead or dying individuals. Species which manifest strong emotional bonds between individuals, including many primates, elephants, and cetaceans, may continue to interact with a corpse for a period of time after the individual’s death. Many such examples involve mothers who continue to carry a dead infant, often for up to a week after death (Goodall 1977; Boesch and Boesch-Achermann 2000; Biro et al. 2010; Fasching et al. 2010), and in some cases other individuals such as unrelated males have been observed to carry or interact with dead infants (Merz 1978). Species with strong social bonds often exhibit emotional responses to dead or dying individuals, ranging from surprise and fear to prolonged grief (King 2013). Cannibalism is also a form of mortuary behavior observed in some non-human primates.

Scientific examination of mortuary behavior across living species has been limited. Piel and Stewart (2015) noted the publication bias related to mortuary behavior in non-human species. Field researchers may hesitate to publish observations that appear to be isolated cases or anecdotes, and as Piel and Stewart point out, almost no studies are negative reports that claim that *no* mortuary behavior occurs in a species. In recent years, a growth in interest in mortuary behavior in nonhuman primates and other social mammals has been fueled by attempts to understand ancestral hominin mortuary practices, including those evidenced in Neanderthals (Pettitt 2018; Pettitt and Anderson 2020; Pomeroy et al. 2020). Discussion of the evolutionary background of mortuary behavior in non-human species has focused upon possible adaptive or maladaptive consequences of the behavior (Piel and Stewart 2015). Less attention has been given to the proximate neural, psychological, or social mechanisms that give rise to mortuary behavior in various species.

There are clear differences between the mortuary behavior repertoire of humans and the mortuary behavior observed in many other living primates. In humans some mortuary behavior is *ritualized*, meaning that the behavior is systematized and repeated. *Funerary behavior* is a category of mortuary behavior which is defined as specific activities relating to the disposal of the dead and to their subsequent commemoration (Pettitt 2011; 2018; Pettitt and Anderson 2020). While funerary behavior is not characteristic of other primates or social mammals, human mortuary behaviors do include many patterns that are also seen in some non-human species, such as the temporary curation of corpses or body parts, disposal of corpses without commemoration, dietary or starvation cannibalism, and manipulation of corpses out of curiosity or fear (Engelke 2019; Silverman et al. 2021).

To the extent that human behaviors surrounding death are different from non-human primates, those differences reflect primarily two mechanisms. One dynamic underlying human mortuary behavior is the shared, learned cultural traditions concerning death. These traditions vary extensively across human cultures and may include an understanding of the permanence of death, religious beliefs and practices concerning death, and scientific and practical knowledge about the causes of death. When anthropologists examine funerary practices, they often describe this kind of cultural knowledge (Engelke 2019; Silverman et al. 2021). Another dynamic involved is an evolved emotional cognition including emotional self-awareness and regulation. In human cultures these two dynamics are interconnected: religious rituals help grieving individuals by providing social support for emotional regulation and for processing and resolving relational trauma, for example. Superficially similar instances of mortuary behavior in different cultures or species may sometimes involve different proximate mechanisms. For example, ritual and funerary behavior both involve the learned repetition of behavioral and emotional states (Pettit 2018, Silverman et al. 2021). In effect a shared capacity for culture integrates with complex emotional cognition, involving a depth of emotional bonds, capacities for emotional commitments (which necessarily come alongside grief at loss, and a need to process relational trauma) and complex emotional regulation. Both have been practiced by people with entirely different or even incommensurable cultural traditions concerning death, which differ widely, yet fulfil similar emotional needs. Hence the interpretation of physical evidence for mortuary behavior should consider the range of cultural and cognitive mechanisms that may be at play and how they interact, which may give rise to different equally plausible explanations for the pattern of evidence.

## Rising Star evidence and context

The findings from the Rising Star system strongly support a scenario where members of the *H. naledi* community carried the bodies of dead conspecifics to more than 30 meters below the surface, over more than 80 meters of underground passages in a difficult and dangerous subterranean environment (Berger et al. 2024; Elliott et al. 2021). The available evidence demonstrates some aspects of mortuary behavior manifested by *H. naledi*. Multiple lines of material evidence show that corpses were manipulated, both at the time of death and afterwards. The distribution of skeletal parts across the Dinaledi Subsystem could not have arisen from deposition at a single point of entry to the subsystem with gravity-driven movement of bodies or bones (Berger et al. 2024). Both the spatial arrangements of skeletal material and the form and composition of sediments rule out water flow or mud flow as mechanisms for transport (Dirks et al. 2015; Wiersma et al. 2020; Brophy et al. 2021; Berger et al. 2024). Movement of remains by *Homo naledi* is the best hypothesis for the emplacement of bodies or remains. The rapid emplacement of some bodies into sediment prior to decomposition and continued support by sediment through the process of decomposition, together with evidence of disruption of surrounding sediment layering, all suggest that some bodies were interred within shallow holes and covered before soft tissue decomposition occurred (Berger et al. 2024). Some bodies were manipulated after deposition, as evidenced by the selective reworking of the Puzzle Box area of the Dinaledi Chamber, leading to fragmentation of some skeletal elements, disaggregation of body parts, and commingling of elements from different individuals (Berger et al. 2024). The presence of young children and infants within the sample likely also reflect the manipulation of bodies by other, presumably older, individuals (Berger et al. 2024, Delezene et al. 2023).

Many aspects of the mortuary behavior represented by this evidence remain unclear. The evidence does not show whether postmortem manipulation in the Puzzle Box area was deliberate and commemorative in intent, or whether this manipulation was an incidental result of the introduction of subsequent bodies into the Puzzle Box are or other activities in the Dinaledi chamber. Archaeological collection and excavation of remains across the Dinaledi subsystem has revealed varied dispositions of different individuals. Some of the skeletal remains show no evidence of intentional disruption after deposition, while others underwent marked and selective reworking after initial deposition, and other skeletal parts were isolated in long fissure passages (summarized in Berger at al. 2024). It is unclear whether these differences reflect intentional differences in mortuary behavior, whether they reflect changing traditions over time, or whether they resulted from a lack of precise patterning of mortuary activity. The duration of mortuary activity in the cave system is not known, nor is it known whether the remains in the Lesedi Chamber and in the Dinaledi Subsystem represent activities of the same group, culturally related groups, or unrelated groups. The anatomical evidence also suggests that the sample of *H. naledi* individuals may be biased with overrepresentation of one sex (Delezene et al. 2024), and it is not known whether this bias was an intentional result of the mortuary behavior. While a majority of the remains associated with mortuary behavior of *H. naledi* occur within deep cave areas, it is not clear whether this behavior was limited to these spaces or whether the observed evidence may represent only a small part of a more extensive pattern.

The use of deep cave spaces as part of the mortuary behavior of *H. naledi* provides additional evidence about the social and emotional mechanisms of this species. The subterranean environment used by *H. naledi* is physically challenging for today’s researchers. A longstanding question is whether the system was equally challenging for *H. naledi*. The journey to the Dinaledi subsystem from any known or reasonably hypothesized incursion point involved strenuous scaling and navigation of complex three-dimensional topography across distance, multiple chambers, passages, climbs and descents (Elliott et al. 2021, Robbins et al. 2021, Berger et al. 2024:SOM). To accomplish this *H. naledi* had to coordinate their behavior and collaborate to move the bodies to a specific location inside the Rising Star cave system. Several aspects of the biology of *H. naledi* suggest that this species may have been better at underground movement than today’s humans: Adult *Homo naledi* individuals had smaller body size than even small-bodied caving team members today (Garvin et al, 2017), and a body plan that was more suited for climbing and passing through narrow and restricted cave passages (e.g. Kivell et al, 2015; Feuerriegel et al, 2017, 2019; Williams et al, 2017; Traynor et al, 2022). Still, movement into these spaces would have had high energetic costs and carried some risk for the *H. naledi* individuals undertaking the behavior. Doing so while carrying a corpse would have entailed additional energetic costs. Given the structural complexity of the cave system layout (Berger et al. 2024, Elliott et al. 2021, Robbins et al. 2021), there must have been some form of explicit communication (tactile, vocal, and likely visual) for coordination of movement and actions, and the potential use of fire as a light source, between the *H. naledi* undertaking the behavior. Such coordination and specific set of actions around the treatment of deceased conspecifics is more methodologically extensive, energetically costly, with higher risk of injury than any reported for other primates and non-human animals to date (King 2013). This behavior is also more complex and multifactorial than that reported for the one earlier case of hominin mortuary behavior (Sima de Los Huesos, Carbonell and Mosquera 2006). There are no clear direct fitness benefits nor any indication of particular proximate functional stimuli for this suite of behaviors.

The subterranean environment used by *H. naledi* is not only physically challenging but is also emotionally and physiologically challenging, reflecting a particular engagement with difficult underground spaces not common in the archaeological record of that time. Dark enclosed spaces, where visual perception is curtailed, can create a state of emotional arousal profoundly affecting perceptual, cognitive, physiological, and social systems (Zuccarelli et al. 2019; Kedar et al. 2021), even with some form of illumination. It is very likely that *H. naledi* used forms of tactile communication and a range of proprioceptive tactics to navigate and communicate in the Rising Star cave system (e.g Dunbar 2010, Waller and Hodgson 2013). However, while such modes of social interactions can enable a range of coordination, it is likely that some form of illumination as also necessary to undertake the behavioral patterns reported (e.g. Pettitt, 2022).

In humans, and other diurnal primates, sensory deprivation through reduced or a lack of consistent visual clues creates a heightened sensitivity to other senses (and thus augments proprioceptive dynamics) as well as prompting experiences of visual disturbances, hallucinations, and disorientation (Hodgson 2021). Experiences of these types of extreme and unusual environments, though often inducing fear responses, can also facilitate powerful bonding experiences (Steidle, Hanke, and Werth 2013). Furthermore, the interoceptive nature of bodily awareness such as that experienced during traversing a complex cave system enhances empathy (Ernst et al. 2013) which may be augmented and deployed in tactile and vocal communication and coordination. This range of substantive emotional, psychological, and physiological reactions may explain why experiences in deep dark caves are often associated with a sense of the transcendent in contemporary humans (Montello and Moyes 2012, Kedar et al. 2021) and given the broad range of sensory commonalities across diurnal anthropoids, and especially apes, such experiences likely had comparable impacts on *H. naledi* and other Pleistocene hominins using subterranean spaces. We argue that careful and coordinated treatment of the dead on several occasions, in these subterranean environments, implies particularly strong social and emotional bonds and some shared understanding of meaning (Pettitt and Anderson 2020) in the handing of the dead by *H. naledi*.

As with most examples of Pleistocene mortuary behavior, researchers should be cautious when comparing to modern analogs. Mortuary and funerary behaviors in the past need not map directly to the practices of contemporary or Late Pleistocene humans (see Berger et al. 2024). As recently suggested by Meneganzin and Killin (2024: 26) in a discussion of Neanderthal aesthetics: “…we should not be surprised if the search for Neanderthal aesthetic practices suggestive of an aesthetic sense requires taking a different route, at least sometimes, to the search for (paradigmatic examples of) human aesthetic practices.” This suggestion is likely applicable to a range of Pleistocene hominin behavior. An important point of comparison is the Neanderthal use of deep caves, in certain cases reflecting a substantial duration of activity and repeated use (Jaubert et al. 2016; Baquedano et al. 2023).

That this high-risk, high-cost, no-overt-direct-fitness-benefit behavior was undertaken repeatedly by multiple members of an *H. naledi* community indicates a valued, likely cultural, tradition with a social and emotional function. The combination of features in the behavior and the context in which it was undertaken (in deep caves with the likely use of fire for illumination), suggests a level of cognitive/semiotic meaning-making capacity in *H. naledi* (e.g., Kissel and Fuentes 2017; 2018; 2021) that matches some similar assessments of other populations of the genus *Homo* during earlier, the same, and later time periods (**Figure 1**). It is our hypothesis that the repetition of mortuary activities within the Rising Star cave system reflects a form of shared memorialization. This hypothesis emerges from the fact that shared attention and joint action were necessary to generate the evidence within the system, and these actions were repeated over some period of time. The hypothesis of shared memorialization does not depend upon individual interments being repeated with identical steps. What supports the hypothesis is the repeated pattern of cave use and the context and distribution of the remains. The collective practice by *H. naledi,* coupled with social and emotional investment, helped transform the ‘space’ of the Dinaledi Chamber and Hill Antechamber to ‘place’ (Low and Lawrence-Zúñiga 2003) through the pattern of mortuary and possible funerary behavior (e.g. Silverman 2008). The coordinated treatment of the dead on several occasions, in these subterranean environments, implies particularly strong social and emotional bonds and some shared understanding of meaning (Pettitt and Anderson 2020) in the handing of the dead by *H. naledi*.

Some form of mortuary behavior by *H. naledi* is a supported hypothesis (Berger et al. 2024). However, there is substantive criticism of the assertions of cultural burials in the Dinaledi subsystem (see Berger et al. 2024, Martinon-Torres et al. 2023, Foecke et al. 2024). Recently Pettitt (2022) laid out the three key criteria for assessing whether or not the *Homo naledi* remains in the Dinaledi subsystem represent actual funerary behavior: a) is there an as-yet unmapped entrance into the Dinaledi Chamber? b) Is there any evidence of artificial lighting in the cave system, and c) Is there evidence that it was dead bodies, rather than body parts that were carried into the chamber? The first query has been repeatedly addressed and no other options for alternative ingress into the Dinaledi Subsystem involving movement across less than 80 meters of structurally complex crevices, chambers, and passages involving a descent to a depth of ∼30 meters below the surface have been found or potentially identified (Elliott et al. 2021, Robbins et al. 2021; Berger et al. 2024). There is reported evidence of fire use (hearths and charcoal and smoke scarring of surfaces) in the Dinaledi system (see Bower 2022, but also see Martinon-Torres et al. 2023) although the age of these occurrences has not been reported yet, so its association with *H. naledi* is currently correlational. However, the correlational support is a reasonable hypothesis given there is no evidence of any kind across the last decade of investigation of any habitual activity by ancient or recent *Homo sapiens* in the Dinaledi Subsystem or even deep proximal spaces such as the Dragon’s Back Chamber or Lesedi Chamber. Articulated skeletal elements do make up a substantive percentage of the *H. naledi* remains reported in the Dinaledi system, especially in feature 1 and in the Hill Antechamber feature (Berger et al. 2024). While the one other major locus of remains in Dinaledi (the Puzzle Box feature and surface collections) have a majority non-articulated materials, there are articulated remains present and there is strong evidence that disarticulation resulted from postdepositional reworking, likely by *H. naledi* (Berger et al. 2024). Therefore, we suggest that the currently available evidence tentatively meets Pettit’s (2022) criterion for funerary action. As it stands, the whole of the evidence supports the hypothesis that the *H. naledi* remains in the Dinaledi subsystem are one of the two earliest examples of a mortuary practice in a hominin, and potentially offer the earliest evidence of multiple interments, postdepositional reworking, and thus funerary actions by a hominin.

In addition to the above, the locations, contexts, and the inferred behavior associated with the *H. naledi* remains most likely demonstrate shared meaning-making activity (Kissel and Fuentes 2017, 2018). Certainly, if the remains are postdepositionally re-worked, and/or do represent burials, and one accepts the correlational association, and validity, of engravings near the interment sites with *H. naledi* (e.g. Berger et al. 2023) they do. But even if one *only* accepts the transport to and placement of bodies in the Dinaledi Chamber and Hill Antechamber locations (and the Lesedi Chamber, Hawks et al. 2017, Berger et al. 2024) there remains a robust argument for mortuary behavior and the assignation of shared meaning to it. Most documented mortuary and funerary practices have been attributed to *Homo sapiens* and Neanderthals, and aside from the Sima de los Huesos site (Carbonell and Mosquera 2006) and Dinaledi, most such evidence is later the Pleistocene (**Figure 1 and Table 1).** Evidence of funerary behavior is generally assumed to require human-like cognitive capability (Pettitt 2018). If such behavior is indeed present in a small-brained hominin it suggests that increases in brain size/EQ are not a necessary precursor for the appearance of complex meaning-making behavior in hominins.

**Table 1.**
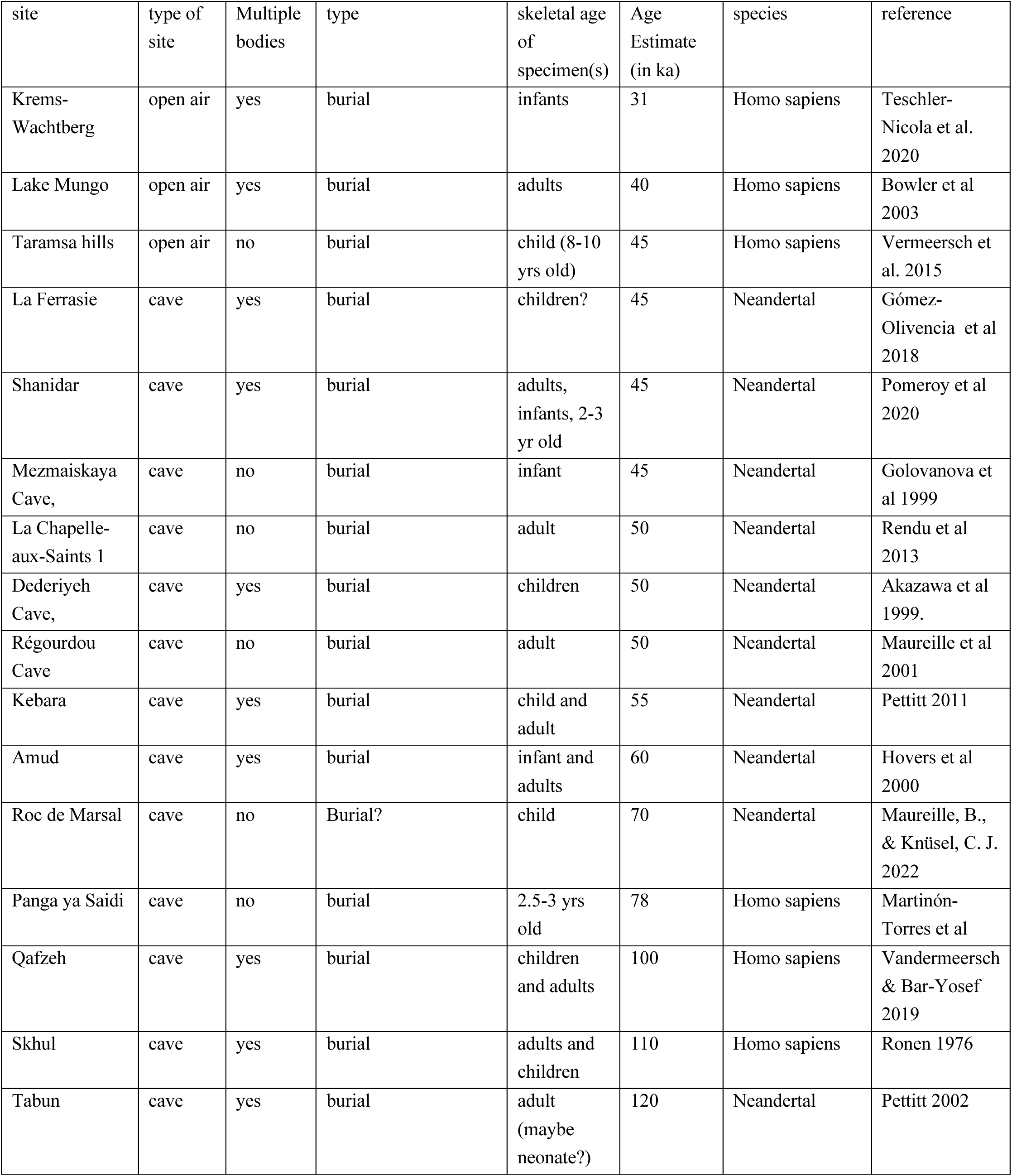
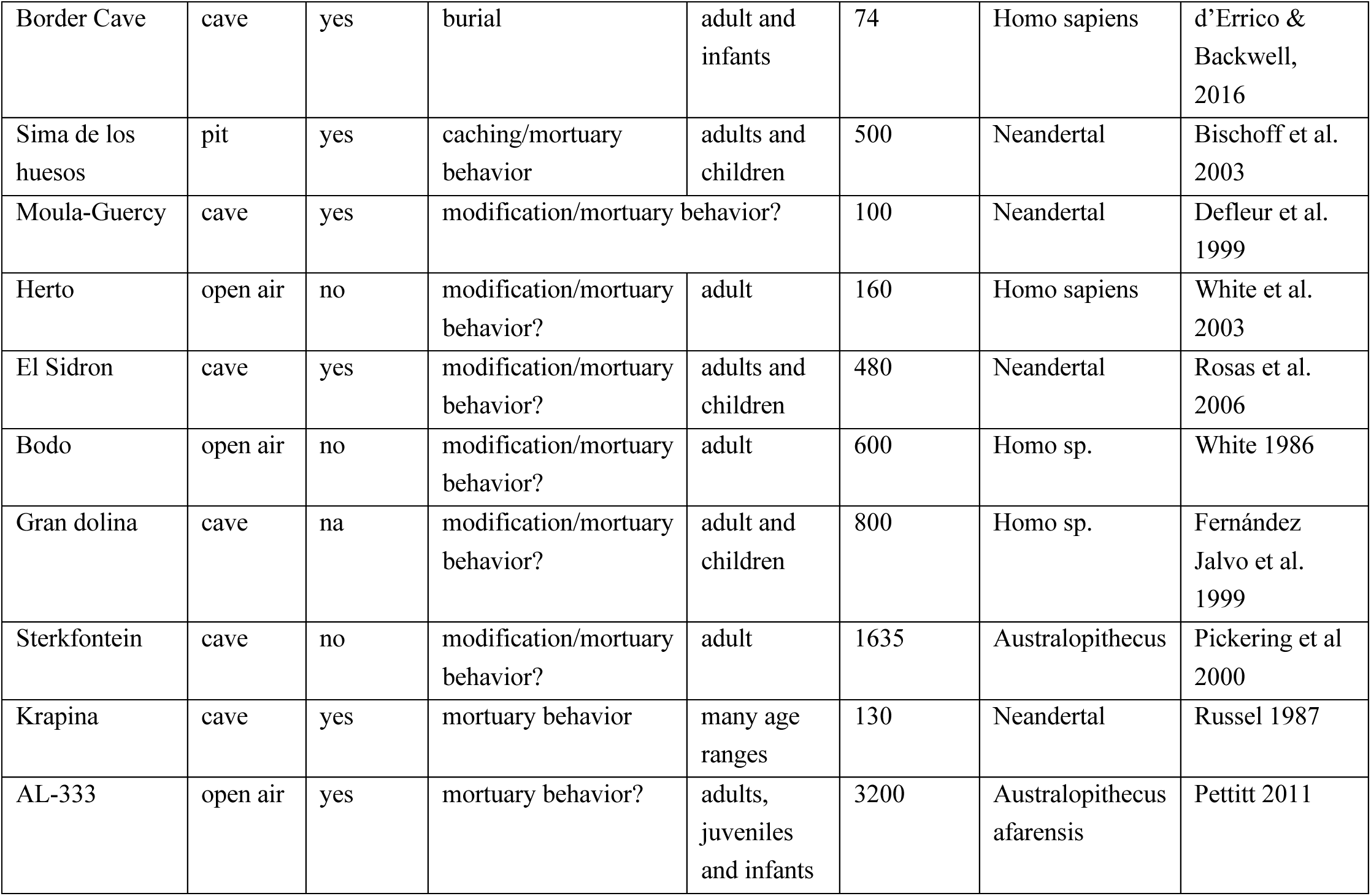
Table of evidence of potential mortuary behavior in hominins. See supplemental material for references

## A role for emotional cognition?

The achievement of social collaboration and social solidarity in humans relies upon emotional cognition and emotional regulation. A broad array data supports the hypothesis that emotional regulation and self-awareness were prerequisites of human social behaviors involving solidarity and cooperation, including cultural learning, language, and provision of extended care (Spikins et al. 2018).

Humans share many building blocks of emotional cognition with other mammals, and some complex abilities with other primates. We share the same visceromotor and sensorimotor foundation for emotions with other mammals for example (Steklis and Lane 2013). Moreover, a range of common emotional responses in humans have now been documented in other primates, particularly in apes, through measurement of heart rate and skin conductance as well as more recently pupil mimicry and infrared thermography (Nieuwburg, Ploeger, and Kret 2021). Interpersonal emotional interactions have a common basis. Emotional contagion is apparent in monkeys and apes, and apes in particular demonstrate a level of empathy though yawning and even sympathy through active consolation (Romero, Castellanos, and de Waal 2010; Preston and de Waal 2002). Diverse primate species have the cognitive ability to infer emotional meaning from expressions (Nieuwburg, Ploeger, and Kret 2021). Moreover, there is anecdotal evidence for the foundations of cognitive empathy in targeted helping within apes (Koski and Sterck 2010).

One important aspect of human emotional cognition is the ability to regulate emotions by bringing feelings into “rational” thought (Green and Spikins 2020). Humans communicate and engage in shared intentions, and meaning-making, to a degree not seen in other animals, and demonstrate motivations to share emotions, experiences and activities with other persons (Steklis and Lane 2013; Fuentes 2018). Emotional self-awareness, and with it the capacity to regulate emotions—that is, to calmly tolerate difficult feelings and bring them into rational thought—is key to many human social behaviors. Emotional self-awareness is essential to translating empathy into systematic compassionate helping for others for example (Spikins 2022). Furthermore, emotional self-awareness allows the regulation of interpersonal emotional vulnerabilities which foster connection and collaboration such as fearfulness (Grossmann 2022). Self-conscious emotions also more broadly regulate social behaviors in general (Beer et al. 2003) and conversely impaired emotional awareness interferes with normal social function in both clinical and nonclinical populations (Gu et al. 2013). Perhaps most importantly emotional self-awareness provides the basis for emotional commitments which bring high levels of give and take to social relationships. Capacities to make emotional commitments to group interests are often demonstrated through costly signaling in risky, nonfunctional ways (Hall et al. 2015; Lang and Kundt 2023).

Another important aspect of human emotional cognition is the presence of emotional cues that are absent or not well developed in other living species. The depth of human emotional commitments come with costs. Firstly, living humans accentuate emotions associated with social control, including shame and guilt, which are not manifested in similar ways in other great apes (Boehm 2012; Turner 2014). Sympathy for others and guilt both emerge early in human ontogeny, with sympathy providing a basis for prosocial orientation in very young children and guilt helping provide motivation for repair of social ruptures (Vaish and Grossmann 2022). These have sometimes been examined within the framework of *moral emotions*, which would additionally include emotions such as contempt, gratitude, and disgust (Fitouchi et al. 2023). These aspects of emotional cognition in humans provide a foundation for prosociality and social solidarity. Both emotional self-awareness and moral emotions function to regulate social interactions and maintain cohesive and cooperative social relationships. Secondly, emotional bonds with high degrees of give and take, and emotional commitments, which drive behaviors such as risky hunting or caring behaviors necessarily involve relational trauma at loss, regulated and resolved through social relationships and cultural practices.

The *H. naledi* evidence suggests that a human-style conscious emotional awareness was present in this hominin despite its small brain size. The hominins carrying out these mortuary activities would need to be able to bring their emotions into “rational thought” in order to both be aware of their own grief and communicate and coordinate shared intentions over the bodies of the deceased. Moreover, they would need to be able regulate their emotions (hold feelings in calm awareness) such that they were able to mutually engage in coordination to carefully negotiate extensive, complex subterranean landscapes, despite the risk and complexity of such behavior, to transport bodies into the Dinaledi Subsystem. This level of emotional regulation, coordination, and awareness is markedly different from the generally personal (such as corpse interaction), sometimes numb (corpse avoidance) and often disordered displays of grief in our nearest relatives (Pettitt and Anderson 2019). The shared and planned transportation and placement of several bodies in the Rising Star system is also evidence of a shared sets of cognitive commitments, beliefs, or assumptions about meaning and action, something similar to what one would term “shared grief” and/or “shared belief” in contemporary humans. The behavioral sequences required for mortuary action also suggests a form of shared memorialization, or at least more behaviorally and communicationally complicated shared attention and action to achieve the deposition of the bodies in the locations in the Rising Star system. Regardless of whether one accepts the interpretation of burials and the presence and association of engravings with *H. naledi*, the underlying cognitive processes associated with just the transport and placement of *H. naledi* into the Dinaledi subsystem indicates a level of conscious emotional awareness that enables and is associated with extensive shared intentionality, forward planning, and repeated cultural behavior involving bodily risk. Equally complex use of caves by Neanderthals (Jaubert et al. 2016; Baquedano et al. 2023) demonstrate a similar emotional self-awareness, and production of highly symmetrical stone tools is also potentially indicative of certain aspects of emotional awareness and regulation in earlier member of the genus *Homo* (Green and Spikins 2020). Furthermore, this evidence suggests a depth of emotional commitments, with a willingness to take risks and costs on another’s behalf. Social understanding of emotions is widely accepted as adaptive in an evolutionary context (Nieuwburg, Ploeger, and Kret 2021) and emotional awareness is associated with better life outcomes in contemporary human contexts (Smith et al. 2023).

That complex emotional cognition is not unique to *Homo sapiens* should not be surprising. But it is not strictly associated with overall brain size. The fact that a small-brained hominin displays these sorts of behaviors suggests that the neurological capacity enabled by a larger than 1000cc brain cannot be the only factor, or necessarily the main factor, enabling the kind of emotional cognition that is considered a central factor in human evolutionary success. Particular brain areas are related to emotional regulation including anterior cingulate cortex (Giuliani et al. 2011) and amygdala (Davidson et al. 2007). Much in the prefrontal cortex has also been implicated in emotional regulation and executive control (Davidson et al. 2007). Recent approaches also focus on large-scale brain networks being implicated in emotional regulation (Morawetz et al. 2020; Pozzi et al. 2021; Rieck et al. 2024). The associations between brain areas or networks and emotional regulation and self-awareness are studied by considering how these cognitive traits correlate with the volumes of various brain areas, and by the activation of brain areas as indicated by oxygen consumption during experimental tasks. Neither kind of study can be compared easily with studies of fossil hominins. The only data on brain anatomy from fossil hominins are the volume of the endocranial cavity and the few sulcal and gyral patterns that imprint on the endocranial surface. These data provide one suggestive indication that emotional regulation may have been important across *Homo*, which is that species of the genus have humanlike frontal cortex configurations, including *Homo floresiensis* and *Homo naledi* (Falk et al. 2005; Holloway et al. 2018; Hurst et al. 2024).

The anatomical data alone do not answer when humanlike emotional self-awareness and emotional regulation first evolved. Some have suggested that hominins would have been under selection for these traits early in their evolutionary history because of the need for cooperating, cohesive groups in open habitats with high predation (Turner 2014). Others have suggested that such behavior is a hallmark of the genus *Homo*, or that abilities such as technical learning or the routine use of fire (Twomey 2019) could only have been manifested in species with emotional cognition that was similar in ways to recent humans. Spikins and Green (2020) specifically pointed to several aspects of Early Stone Age artifacts like Acheulean handaxes as possible indicators of self-control including inhibition and conscious regulation of emotions. The reported *H. naledi* behavioral activities, may have depended on emotional commitments to others combined with a set of cultural beliefs/practices, a high level of emotional awareness to manage these, and in turn collaboration with extensive coordination.

## Homo naledi behavior in broader perspective

The behavior patterns manifested in the Rising Star cave system have a distinctive place within a broader global context of mortuary behavior. The repeated mortuary behavior involving more than 30 individuals from Sima de los Huesos, Spain, is substantially earlier than evidence from the Dinaledi Subsystem (Carbonell and Mosquera 2006; Arsuaga et al. 2014; Vives 2015; Sala et al. 2024). Cutmarks on hominin individuals such as the Bodo 1 skull from Bodo, Ethiopia, and the StW 53 skull from Sterkfontein, South Africa, may result from even earlier mortuary activity by hominins (White 1986; Pickering et al. 2000). In the case of StW 53, the evidence is associated with a skull that many researchers attribute to *Australopithecus,* although the claim of hominin-produced cutmarks has been disputed (Hanon et al. 2018). Curation of hominin skeletal remains was part of mortuary behavior at Herto, Ethiopia before 160,000 years ago (White et al. 2003; Clark et al. 2003). Archaeological evidence of mortuary behavior becomes increasingly common in later contexts, and within the Late Pleistocene, burials and other kinds of funerary behavior have been attributed to both modern humans and Neanderthals. Within this broader pattern, the Rising Star evidence stands out in two ways: the early possible occurrence of burials (Berger et al. 2024; Dirks et al. 2017), and the position of *H. naledi* as a phylogenetic outgroup when compared to modern humans and Neanderthals (Dembo et al. 2016; Argue et al. 2017; Caparros and Prat 2021).

Mortuary behavior is only one category within a broader set of behavior patterns that emerge from shared meaning within social groups **(Table 1 and Figure 1)**. Another category of evidence is the patterned engraving of lines on bones, shells, rocks, or rock walls. Providing geochronological context for such engraved features on rock walls is challenging as it requires that walls themselves be buried in sediment or that engraved features be partially obscured by material susceptible to dating, such as calcite crusts. The engraved lines and percussion marks observed within the Dinaledi Subsystem (Berger et al. 2023) are not an exception to this challenge; however, the widespread presence of *H. naledi* remains within this space and absence of any evidence of modern human activity other than entry by recent explorers, make it a reasonable hypothesis that *H. naledi* individuals were authors of these marks. Such a hypothesis is reasonable when considered in a global context. Engraved bones are known from several sites as old or older than the Dinaledi Subsystem, including marked bones, ivory, and stone from Bilzingsleben, Germany (Mania and Mania 1988), an engraved shell from Trinil, Indonesia (Joordens et al. 2016), and an engraved bone from Kozarnika, Bulgaria, once suggested to be around 1.4 million years old (Guadelli and Guadelli 2004), although recent geochronological work suggests that the layer is likely early Middle Pleistocene in age (Heydari et al. 2022). A larger array of such evidence is known from later Middle Pleistocene and Late Pleistocene, including examples of engraved hominin bone.

Evidence of the entry into caves by hominins occurs across the span of the fossil record of South Africa, three million years or earlier. What is apparently somewhat unusual in the Rising Star Cave is the repeated hominin use of deep caves, where illumination is necessary, including nonutilitarian use of the space. No evidence of such behavior has yet been found earlier than the Dinaledi Subsystem. Still, the evidence for manipulation of stalagmites and built structures within Bruniquel Cave, France (Jaubert et al. 2016), presumed to have been done by Neanderthals, as well as later Neanderthal marking within deep caves such as Cueva de Ardales, Spain (Pitarch Martí et al. 2021), or areas far from cave entrances such as at Gorham’s Cave, Gibraltar (Rodríguez-Vidal et al. 2014), bring to mind the activity seen in the Rising Star cave system. Within South Africa, the site of Wonderwerk Cave has evidence that may be contemporary or slightly later than the Dinaledi Subsystem including ochre and quartz crystals taken approximately 100 m from the cave entrance, although within this linear and large cave the entrance is always visible from the excavated area (Chazan et al. 2020).

The sustained use of deep cave areas is, in humans, tied to the use of fire for illumination. Hominin control of fire in South Africa long predates the Rising Star evidence, with controlled fire indicated in Member 3 at Swartkrans (only ∼ 800m from the Rising Star Cave), and in Early Stone Age context at Wonderwerk Cave, by one million years ago (Brain and Sillen, 1988; Brain, 1993; Berna et al., 2012). Both *Paranthropus* and early *Homo* were present in South Africa during that period and occur in association with combustion evidence (Brain and Sillen, 1988; Brain, 1993). Additionally, cave use by hominins across multiple locations during the period of *H. naledi* and earlier (e.g. Wonderwerk in South Africa and Bruniquel in France) also involved use of fire (Berna et al. 2012, Jaubert et al. 2016). While we cannot yet be certain of the exact modes, intensity, duration, and quality of the fires used by *H. naledi* in the Dinaledi subsystem, it is a strong assumption that they at least provided flickering and moderate intensity light sources.

Use of ochre and other pigments is another category of behavior linked to meaning-making (Dapschauskas et al. 2022; Kissel and Fuentes 2018). Material evidence of pigments carried or used by hominins has been found at several sites earlier than the Dinaledi Subsystem, including sites within Africa, as well as within southwest Asia and south Asia **(Figure 1)**. Non-utilitarian objects that were transported by hominins into sites include some with physical or iconic resemblance to human figures, including objects from Berekhat Ram, Israel (Goren-Inbar 1986) and Tan-Tan, Morocco (Bednarik 2003), both from the later Middle Pleistocene.

This pattern of evidence shows that *H. naledi* and other populations of the genus *Homo* overlap temporally in the expression of meaning-making behavior. The material evidence indicates some degree of shared socioemotional and cognitive processes. Middle Pleistocene hominins varied in brain sizes and cranial and post-cranial morphologies, and many of these varied populations share increased evidence for meaning-making (**Figure 1**). Such behavior is neither “modern” nor exclusive to larger brained *Homo sapiens* (and Neanderthals). Whilst this adds further evidence to our understanding of the emergence of hominin cognition there are also wider evolutionary implications. Much like the evolution of social emotional abilities in other primates (Nieuwburg, Ploeger, and Kret 2021) the behavioral evidence for small-brained *H. naledi* may suggest that some degree of analogous, and homologous, evolution underlies social emotional complexity in humans.

## Reconsidering brain size

Since the nineteenth century, ideas about the evolution of human behavior have tended to emphasize that larger brains evolved to enable more complex behavior. In the most general sense this is surely correct. Large brains are expensive to maintain and grow (Isler and van Schaik 2009). Considering these costs, the large brain sizes manifested in some hominin lineages would not have evolved if they were not reliably correlated with survival or reproduction. It is often assumed that a large brain was an essential step towards a uniquely human cognition, social relationships and culture (Dunbar 2003; Muthukrishna et al. 2018). Several hominin lineages during the Pleistocene did experience evolution of larger overall and relative brain size, as measured by endocranial volume **(Figure 2)** and body size. Nothing in the Pleistocene behavioral record refutes the basic idea that such lineage-specific increases in brain size evolved alongside some behavioral adaptations.

But the Pleistocene record today does not support the notion that *every* behavioral adaptation was mediated by overall brain size. Planning and forethought in stone tool production both preceded the first appearance of *Homo* (Harmand et al. 2015). Early populations of *H. erectus* dispersed from Africa into Eurasia before 1.8 million years ago, and the first substantial sample of these hominins has endocranial volume ranging from 550 ml to 730 ml (Ponce de Léon et al. 2022). The first bifacial tool traditions were developed before 1.76 million years ago, within a geographic and temporal context where the only extant groups of hominins had small average endocranial volumes (Lepre et al. 2011). The use of fire emerged in excess of a million years ago in Africa (Hlubik et al. 2019) at a time when the average endocranial volume among crania attributed to *H. erectus* was around 750 ml. It is credible that ancestors of *H. naledi* were among these early fire users. The phylogenetic arrangements of species within the genus *Homo* remain uncertain (Hawks et al. 2017; Argue et al. 2017; Dembo et al. 2016; Caparros and Prat 2022). Still, much evidence suggests that the lineages leading to hominin species with small brain size including *H. naledi* and *H. floresiensis* emerged from within this early Pleistocene evolutionary context (Dembo et al. 2016; Hawks and Berger 2019). These species would have been part of the hominin niche that also gave rise to other later Pleistocene members of the genus *Homo* (Mondanaro et al. 2020).

It is also important to be precise about what the small sample of fossils really tells us about differences in brain size between lineages (Figure 2). Known *Homo naledi* individuals for which endocranial volume estimates can be made (n = 5) have volumes ranging from 450 ml to 610 ml (Holloway et al. 2018; Hurst et al. 2024), the single known endocranial volume for *H. floresiensis* is 426 ml (Kubo et al. 2013; Falk et al. 2005). With so few fossils, these samples are unlikely to provide accurate understanding of the average of either species. Moreover, both samples may be biased. The LB1 individual of *H. floresiensis* is inferred to be female based on postcranial morphology (Brown et al. 2004) and male individuals would likely have had larger endocranial volumes. The low degree of variation in size within the *H. naledi* sample suggests that the sample may be biased toward representation of one sex (Delezene et al. 2024), again indicating that the sample average may misrepresent the species mean. Both samples have smaller average endocranial volume than *H. erectus*, yet many the fossils attributed to this species within Africa have estimated endocranial volumes within or just above the range observed in *H. naledi*, including DNH 134 (Herries et al. 2020), KNM-ER 42700 (Neubauer et al. 2018), DAN5/P1 (Semaw et al. 2020), KNM-OL 45500 (Potts et al. 2004), and OH 12. While global *H. erectus* endocranial volume increased over time (Leigh 1992), fossils attributed to this species with small endocranial volumes occur across the entire temporal range of the species within Africa.

Different hominin lineages manifested differences in brain organization, and these do not always correspond to changes in brain size (Holloway et al. 2018; Hurst et al. 2024; Ponce de Léon et al. 2021). Endocast evidence from across the genus *Homo* suggests that humanlike frontal cortex development and morphology occurs across a broader array of species than large brain size (Falk et al. 2005; Holloway et al. 2018; Ponce de Léon et al. 2021). *Homo floresiensis* and *Homo naledi* both share some humanlike aspects of frontal lobe organization while they both have brain sizes within the range of *Australopithecus* (Falk et al. 2005; Holloway et al. 2018; Hurst et al. 2024). Early *Homo erectus* is argued to be polymorphic in frontal lobe organization (Ponce de Léon et al. 2021), and as noted above many fossil crania attributed to this species have endocranial volumes smaller than 650 ml (Lordkipanidze et al. 2013; Ponce de Léon et al. 2021; Herries et al. 2021; Semaw et al. 2020).

## Looking forward

The initial rise of complex social behaviors in hominins may have been fueled by evolution of emotional cognition alongside other cognitive processes. Meaning-making by hominins includes a variety of categories of behavior that are joined together by the role of shared intention, repetition, and social collaboration. Mortuary behavior is one of these. Varied expressions of mortuary behavior are present in other social mammals, and hominins are distinct in the extent of repetition, social collaboration, and shared meaning in the process. Rather than relying on increased encephalization and its relation to complex behavior as a *Cognitive Rubicon* in human evolution (see Meneganzin and Currie 2022), we suggest that a distinctive cultural, empathetic, collaborative niche dependent on increasingly complex and robust relationships between individuals has also been a primary driver in the development of key aspects of human, or human-like, behavior (Galway-Witham, Cole, and Stringer 2019; Kissel and Fuentes 2021; McBrearty and Brooks 2000; Fuentes 2017; Spikins 2022; DeCasien, Barton, and Higham 2022). The increasing data for complex behavior and meaning-making across the Pleistocene should play a major element in structuring how we investigate, explain, and model the origins and patterns of hominin and human evolution (Kissel and Fuentes 2021; Spikins 2022; Spikins et al. 2019). The current evidence for *H. naledi* in the Rising Star system pushes back the origins of mortuary, and possibly funerary behaviors, challenges our assumptions about the role and importance of encephalization in human evolution, and suggests that the hominin emotional, socio-cognitive niche is more significant than previously thought.

## Acknowledgments

Permits to conduct research in the Rising Star Cave system are provided by the South African National Research Foundation (LRB). Permission to work in the Rising Star cave is given by the LRB Foundation for Research and Exploration. The Authors would like to acknowledge the funders of the various expeditions and documentation of the engravings including the National Geographic Society (LRB), the Lyda Hill Foundation (LRB), and the National Research Foundation of South Africa (LRB). Laboratory work and travel was funded by the National Geographic Society (LRB), the Lyda Hill Foundation (LRB), the Fulbright Scholar Program (JH), the University of Wisconsin (JH), and Princeton University and the John Templeton Foundation (AF). We also want to express a sincere thanks for the inspiration of the late, great Calvin Keys.

## Author contributions

Conceptualization: AF, LRB, JH, MK, PS

Methodology: AF, MK, LRB, JH

Investigation: AF, PS, MK, LRB, JH, KM

Visualization: MK, JH

Funding acquisition: LRB, AF

Project administration: AF, LRB, JH

Supervision: AF, LRB

Writing – original draft: AF, PS, MK

Writing – review & editing: AF, PS, MK, LRB, JH, KM

## Competing interests

Authors declare that they have no competing interests.

## Data and materials availability

All data, code, and materials used in the analysis are available in the SOM and in Berger et al (2023, 2024)

## Supplementary Materials

references for Table 1 on mortuary practices

supplement for Figure 1

## Supplement for Table 1 on mortuary practices

## Text for Supplement table 1 on mortuary practices

Note on table 1: Table 1 was constructed by searching the literature for examples of funerary behaviors in the Paleolithic. The nature of many of these finds are contentious (1). For example, some scholars have rejected Roc-de-Marsal (2, 3) as a burial based on reevaluating the context of the site, while others include it in list of European burials (). Similarly, experts are divided as to if La Chapelle-aux-Saints should (4) or should not (5, 6) be accepted as an intentional Neandertal burial. We also list sites that are not burials but instead show possible evidence of modifying the body after death. Again, scientists disagree if these are funerary practices or, in the case of cutmarks on hominin bones, cannibalism.

Most archaeologists define burial in a way that lets them detect it archaeologically (excavate a pit, put body in pit, refill pit), but this is only one type of funerary ritual. All cultures must find ways to deal with the beginning of life and with the end of life. How to treat the dead (and how to decide when someone is really dead) is culturally specific. The symbolic practices related to death are also highly varied. Just as with foodways, deathways are mediated by how a culture sees death.

All of this is to say that the symbolic aspects around death are as important as the process of dealing with the body. We might also ask how the death related symbolic behaviors help people mourn.

## Supplementary Material

### Data for Figure 1

#### Table used to create figure 1

**Table X.** Data used to create **Figure 1**.

| site | group | type | Max.age | Min.age | Midrange | associated species | references |
| --- | --- | --- | --- | --- | --- | --- | --- |
| Bruniquel | non-functional cave use | cave | -0.176 | -0.176 | -0.176 | Neandertals | Jaubert et al 2016 |
| Wonderwerk | non-functional cave use | cave | n/a | n/a | -0.18 | unknown archaic | Chazan and Horowitz 2009 |
| Cueva de Ardales | non-functional cave use | cave | n/a | n/a | -0.66 | Neandertals | Martí et al. 2021 |
| Lingjing | engraved objects | engraved bone | -0.125 | -0.105 | -0.115 | archaic Homo | Li et al 2019 |
| Kozarnika | engraved objects | engraved bone | -1.6 | -1.4 | -1.5 | n/a | Sirakov et al. 2010 |
| Trinil | engraved objects | engraved shell | -0.64 | -0.38 | -0.51 | Homo erectus | Joordens et al. 2014 |
| Bilzingsleben | engraved objects | engraved bone | -0.37 | -0.23 | -0.3 | Homo erectus | Mania and Mania 1988 |
| Diepkloof Rock Shelter | engraved objects | engraved eggshell | -0.119 | -0.099 | -0.109 | n/a | Texier et al 2013 |
| Qafzeh | engraved objects | engraved stone | -0.118 | -0.075 | -0.0965 | Homo sapiens | Hovers et al. 1997 |
| Sainte-Anne | engraved objects | engraved | -0.21 | -0.17 | -0.19 | n/a | Raynal and Seguy 1986 |
| Oldisleben | engraved objects | engraved bone | -0.13 | -0.08 | -0.105 | n/a | Bendarik 2006 |
| Sima de los Huesos | mortuary behavior | mortuary | -0.6 | -0.4 | -0.5 | Neandertals | Bischoff et al. 2007 |
| Rising Star Cave | mortuary behavior | mortuary | -0.335 | -0.241 | -0.29 | Homo naledi | Berger et al 2022 |
| Skhul | mortuary behavior | mortuary | -0.13 | -0.1 | -0.115 | Homo sapiens | Ronen 1976 |
| Tabun | mortuary behavior | mortuary | -0.12 | -0.12 | -0.12 | Neandertals | Pettitt, P. B. (2002) |
| Qafzeh | mortuary behavior | mortuary | -0.13 | -0.08 | -0.105 | Homo sapiens | Vandermeersch and Bar-Yosef 2019 |
| Panga ya Saidi | mortuary behavior | mortuary | -0.082 | -0.074 | -0.078 | Homo sapiens | Martínón-Torres et al. 2021 |
| Berekhat Ram | non-utilitarian objects | art | -0.47 | -0.233 | -0.35 | n/a | d'Errico and Nowell 2001 |
| Makapansgat | non-utilitarian objects | art | n/a | n/a | -3 | australopithecus | Bednarik 1998 |
| Tan-Tan | non-utilitarian objects | art | -0.5 | -0.3 | -0.4 | n/a | Bednairk 2003 |
| Krapina | non-utilitarian objects | ? | -0.13 | -0.08 | -0.13 | Neandertals | Radovčić et al 2016 |
| Gadeb 8e | ochre use | ochre | -1.45 | -0.75 | -1.1 | n/a | Clark and Kurashina 1979 |
| Bizat Ruhama | ochre use | ochre | -1.96 | -0.78 | -1.37 | n/a | Ronen et al. (1998) |
| Kathu Pan 1b | ochre use | ochre | -0.417 | -0.417 | -0.417 | n/a | Watts et al. 2016 |
| Hungsi | ochre use | ochre | -1.5 | -0.3 | -0.9 | n/a | Watts et al 2014 |
| Maastricht-Belvedere C | ochre use | ochre | -0.27 | -0.2 | -0.235 | Neandertals | Roebroeks et al 2012 |
| GnJh-15 | ochre use | ochre | -0.285 | -0.285 | -0.285 | n/a | Denio and Mcbrearty 2003 |
| Biddenham | ornamentation | ornamentation | -0.3 | -0.3 | -0.3 | n/a | Bednarik 2005 |
| Contrebandier's Cave | ornamentation | ornamentation | -0.115 | -0.11 | -0.1125 | n/a | d'Errico et al 2009 |
| Krapina | ornamentation | ornamentation | -0.13 | -0.13 | -0.13 | Neandertals | Radovicic et al. 2015 |
| Border Cave | ochre use | ochre | na | na | 0.2 | Homo sapiens | Wadley et al. 2020 |
| Rising Star Cave system | engraved objects | Cave wall | -0.335 | -0.241 | -0.29 | Homo naledi | Berger et al 2023b |

## Text for notes on table for Figure 1.

This table is a sampling of archaeological sites that have been suggested to show signs of what some call “symbolic behavior.” Delimitating what is and what is not symbolic has been the source of contention for many decades now (1–6). Traditionally, archaeologists have defined symbols as objects that have meanings embedded in them. Yet a symbol, by its very nature, must be interpreted within a system of meaning and discerning if something is symbolic becomes difficult without knowing the cultural context within which it has been created (7). We created this table from the published literature to demonstrate that no matter what we choose to call it, culturally-mediated behaviors predate contemporary humans. Such behaviors are found with Homo erectus (8), Neandertals (9) and other archaic populations (10–12).

## Notes

### Competing Interest Statement

The authors have declared no competing interest.

### Summary of Updates

While we took great effort to address all of the comments directed at the contents of this manuscript and improve the argument in the light of feedback, we note at the outset that the majority of critique we received was targeted not at the contents of this preprint but at the two other preprints that this one accompanied ( Evidence for deliberate burial of the dead by Homo naledi and 241,000 to 335,000 Years Old Rock Engravings Made by Homo naledi in the Rising Star Cave system, South Africa). It is not our place here to respond to the specific critiques of the other two manuscripts. The respective groups of authors have done so separately. This process has led to an inevitable delay in finalizing this revision. Considering the critiques of, and debate around, H. naledi burial and the fact that the reviewers saw this paper to rely upon the other data, we waited for the revision of the burial manuscript to be posted before we posted the revised version of this one.In this revised manuscript we clarify the goal, intent, and argument of this paper to maximize effective critical reading of its contents. We appreciate and look forward to continued critique and enhanced discussion of this topic and argument.

